# *Cryptococcus neoformans* adapts to host CO_2_ concentrations through the coordinated remodeling of central carbon metabolism, oxidative stress resistance, and membrane homeostasis

**DOI:** 10.1101/2025.11.28.691145

**Authors:** Laura C. Ristow, Emma E. Blackburn, Andrew J. Jezewski, Xiaorong Lin, Damian J. Krysan

## Abstract

*Cryptococcus neoformans* is an environmental pathogen that remodels its cellular physiology to survive within mammals and, in susceptible hosts, cause life-threatening meningoencephalitis. Of the many distinctions between the external environment and mammalian tissues, CO_2_ concentration in the host is 2 orders of magnitude higher than in the environment and represent a critical stress for *C. neoformans*. *C. neoformans* strains that do not replicate at host CO_2_ concentrations are less virulent in mouse models of infection, further supporting CO_2_ tolerance as a virulence trait. To further understand the genetic determinants of *C. neoformans* CO_2_ tolerance, we performed a near genome-wide screen for deletion mutants with altered CO_2_ fitness using a competitive growth assay. A total of 301 of 4698 deletion mutants showed altered CO_2_ tolerance (245 reduced fitness; 51 increased fitness) demonstrating the global effect of host CO_2_ on *C. neoformans* physiology. Based on this data set as well as a metabolomic analysis of *C. neoformans* adaptation to host CO_2_, we show that remodeling of central carbon metabolism, oxidative stress buffering and membrane homeostasis represent an integrated response to CO_2_ stress that is mediated in part by the TOR-Ypk1 signaling axis. We propose that CO_2_-induced capsule formation leads to reduced cellular glucose which, in turn, triggers remodeling of central carbon metabolism toward utilization of alternative carbon sources and increased mitochondrial respiration/reactive oxygen generation. Thus, these data provide a near genome-wide profile of the genetic determinants *of C. neoformans* CO_2_ tolerance as well as a model for how this important environmental human fungal pathogen alters its physiology to proliferate in the host.

## Introduction

*Cryptococcus neoformans* is an environmental fungus that causes life-threatening infections of the central nervous system in humans (1). People with altered immune function such as organ transplant patients and those living with HIV/AIDS are at particular risk for invasive cryptococcal infection (2). *C. neoformans* has been cultured from a wide-range of environmental niches throughout the world (3). The majority of environmental *C. neoformans* isolates are non-pathogenic in mammalian models of infection whereas *C. neoformans* isolated from patients are generally more virulent (4). One well-established reason for this distinction is that most environmental strains of *C. neoformans* are unable to grow at mammalian body temperature (∼37°C, ref. 5). Growth at human body temperature, polysaccharide capsule formation and cell wall melanization represent three of the most widely studied *C. neoformans* virulence traits (5). Despite the prominence of the “big-three” virulence traits (6), other aspects of *C. neoformans* biology contribute to its ability to cause disease and the identification of traits that can better distinguish pathogenic and non-pathogenic strains remains an area of strong research interest in fungal pathogenesis (7).

In addition to elevated body temperature, the mammalian host environment presents additional stressors that could also represent significant fitness bottlenecks to *C. neoformans*; low levels of nutrients and reactive oxygen species (ROS) are two commonly cited examples (6, 7). Our group found that the elevated concentrations of CO_2_ present in host tissues (5%) relative to the external environment (∼0.04%) represents a significant stressor for *C. neoformans* (8, 9). Overall, environmental strains have reduced fitness in host concentrations of CO_2_ relative to clinical isolates. Importantly, environmental strains that are CO_2_-tolerant are also able to cause disease in mouse models (10). In addition, CO_2_-sensitive environmental strains recovered from infected mice have increased CO_2_-fitness as well as increased fitness in the host (11). Taken together, these data strongly support the concept that tolerance of host CO_2_ concentrations is a virulence trait for *C. neoformans* (9).

We have been interested in identifying the genetic and physiologic determinants of *C. neoformans* CO_2_ tolerance. In previous work, we showed that the target <u>o</u>f rapamycin (TOR) pathway is a critical mediator of CO_2_ tolerance and that, in part, this was due to the role of TOR in remodeling of phospholipid asymmetry of the plasma membrane in response to host CO_2_ (12). In addition, we found that multiple genetic loci contribute to the CO_2_ fitness of tolerant strains through quantitative genetic trait analysis of backcrossed strains (11). To gain a more complete profile of the genes that affect CO_2_ tolerance, we carried out a large-scale competitive fitness screen using the *C. neoformans* deletion mutant libraries constructed by the Madhani lab (13).

The results of this screen have provided new insights into the breadth of cellular systems affected by host CO_2_ as well as the multiple stress response pathways that contribute to the ability of the CO_2_ tolerant reference strain H99 to compensate to elevated, host-relevant concentrations of CO_2_. We focused our follow-up studies on the metabolic and oxidative stress pathways and processes required for CO_2_ tolerance. The results of these experiments suggest that metabolic remodeling, buffering of ROS species, and membrane homeostasis represent an interconnected and integrated response to high CO_2_ concentrations that is critical for *C. neoformans* to adapt to this host-relevant stressor.

## Results

### The transition to host CO_2_ concentrations affects a wide range of cellular processes in *C. neoformans*

To identify genes with altered fitness at host concentrations of CO_2_ (5%), a moderate throughput competitive fitness assay that we previously reported (12) was used to screen 4698 strains from the Madhani *C. neoformans* deletion library; all deletion mutants were in the CO2 tolerant H99 strain background (Table S1). Each mutant was inoculated into the well of a 96-well plate with an approximately equal cell concentration of mNeonGreen-labeled reference strain H99. The strains were incubated in RPMI buffered with MOPS in ambient air or 5% CO_2_ at 37°C for 24hr. The ratios of the two strains were quantified using flow cytometry and the competitive fitness calculated as described in materials and methods. Mutants with statistically significant (three replicates, p <0.05 Student’s t test) competitive fitness changes of ±0.2 were considered to have altered fitness. At total of 301 genes met these criteria with 245 mutants showing reduced fitness and 56 mutants displaying increased fitness (Fig. 1A, Table S1). In addition, multiple kinase mutants that we previously showed have altered fitness in 5% CO_2_ were also in this library and showed the same fitness phenotypes (12), including mutants of *MPK1*, *BCK1*, *MKK2*, *PKA1*, *ARK1*, *PKA1*, *CBK1*, *FBP26*, *YPK1*, and *CEX1*. These data provide strong support that the set of mutants with altered fitness in 5% CO_2_ in tissue culture medium is robust.

**Figure 1:**
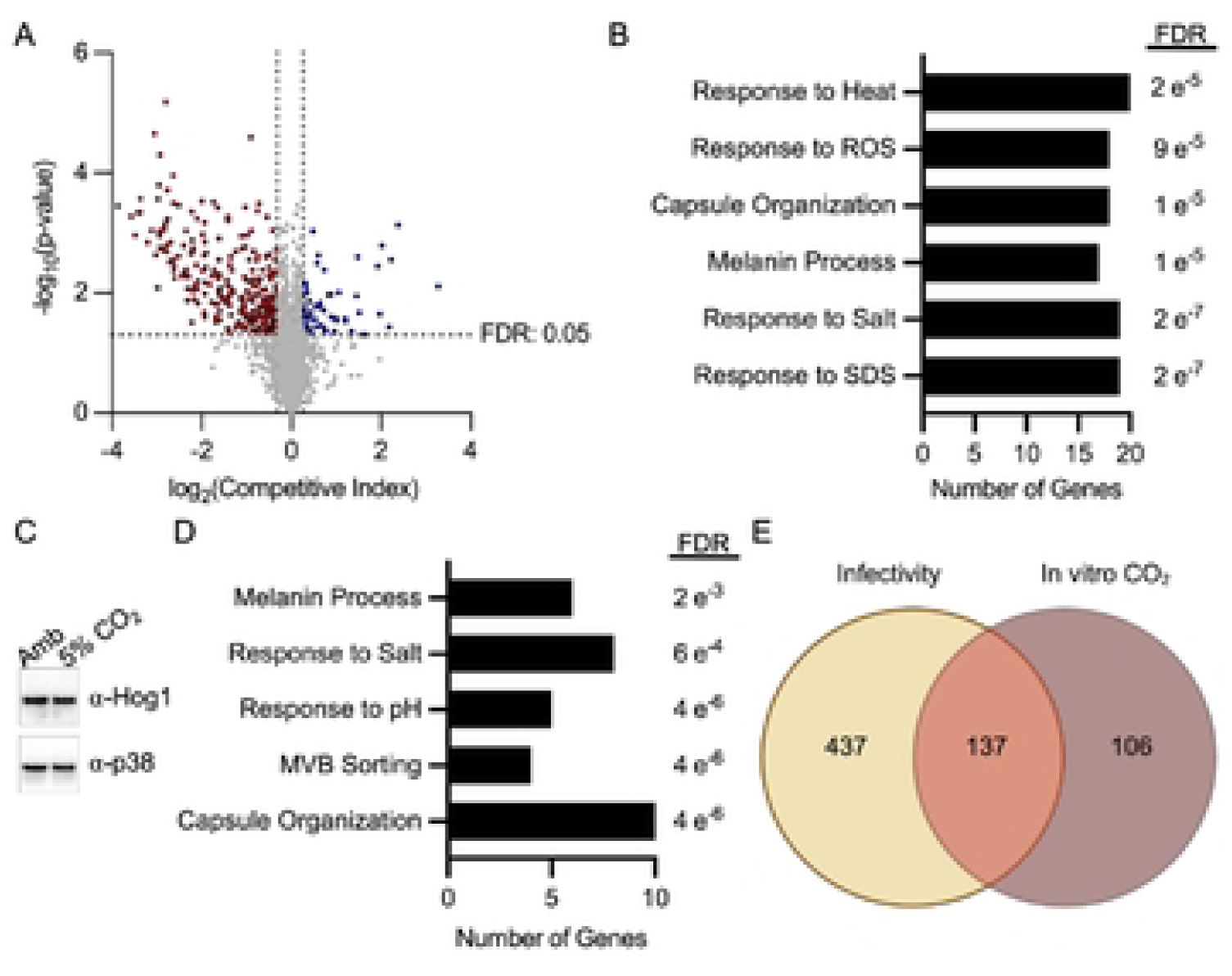
Genome-wide deletion collection screening revealed a wide range of cellular processes involved in response to CO_2_. **A**. Volcano plot representing the average competitive fitness score and significance of three biological replicates for each mutant in 5% CO_2_ conditions relative to ambient air in the Madhani deletion collection. Competitive fitness was determined after combining a 1:1 ratio of mNeonGreen labeled H99 and unlabeled mutant cells in RPMI 1640 medium with 165 mM MOPS, pH 7 and incubating at 37°C in ambient air or 5% CO_2_ for 24 hr. Cell populations were characterized by flow cytometry and the percentage of mNeonGreen negative cells in 5% CO_2_ was divided by those in ambient conditions to calculate competitive fitness. Colored data points indicate values that were below 0.8 (red) or above 1.2 (blue) and statistically significant by Student’s t-test (P < 0.05). Biological-process GO term analysis for the genes identified in (**A**) at 24h are depicted in **B** (significantly reduce CO_2_ fitness) and **D** (significantly increase CO_2_ fitness). **C**. Cell lysates from H99 cells exposed to ambient air or 5% CO_2_ for two hours were analyzed by western blot for Hog1 expression and phosphorylation status. Results are representative of three biological replicates. **E**. Venn diagram comparing mutants with reduced CO_2_ fitness (identified in **A**) to mutants with reduced fitness in a mouse model of cryptococcosis reported in ref. 13.

Biological process GO term analysis of the set with reduced fitness provided some insight into the overall functions involved in CO_2_ tolerance (Fig. 1B). Response to the detergent sodium dodecyl sulfate (SDS) and salt were the two terms with the lowest adjusted p values (Benjamini-Hochberg). SDS interferes with membrane function and is consistent with our previous findings that CO_2_ leads to remodeling of the plasma membrane and that drugs targeting sterols (fluconazole) and sphingolipid biosynthesis (myriocin) are more active against *C. neoformans* at host CO_2_ concentration than in ambient air (8).

The response to salt appears to be driven by a large set of protein kinases with pleomorphic phenotypes (14) that include sensitivity to salt (*CCK1*, *IPK1*, *CBK1*, *YPK1*, *SNF1*, *TPS2*). If the host CO_2_ concentrations directly triggered a stress response that overlaps with salt stress, then it may increase activity of the <u>H</u>yper<u>o</u>smolar <u>G</u>lycerol <u>P</u>athway (HOG) pathway, an important part of the cellular response to elevated salt (14). Our previous screen of a protein kinase library (12), however, did not find that mutants of the *HOG1* pathway were less fit in CO_2_ (in the current screen the *hog1*Δ mutant showed modestly reduced fitness 0.78, Table S1). Furthermore, transition to high CO_2_ did not change the phosphorylation status of Hog1 (Fig. 1C). Therefore, it seems sensitivity to salt is an indirect correlation with CO_2_ sensitivity. Finally, genes involved in response to oxidative stress were also enriched in those required for CO_2_ fitness.

The set of gene mutations that increased CO_2_ fitness is also enriched for response to salt along with response to pH and <u>m</u>ulti<u>v</u>esicular <u>b</u>ody (MVB) formation (Fig. 1D). The presence of the latter two groups of genes is likely related to the fact that the Rim101 pathway is maladaptive during CO_2_ stress (12) and that some genes that contribute to the MVB are involved in Rim101 processing (15). Interestingly, the set of genes required for full CO_2_ fitness was enriched for those that are also annotated to affect the “big three” virulence traits (5, 6) of capsule formation, melanin metabolism and response to heat (Fig. 1B). However, the set of genes whose loss of function increases fitness in 5% CO_2_ was also enriched for those with effects on capsule, melaninization, and salt stress (Fig. 1D). The connection between genes that affect CO_2_ fitness and the “big three” virulence traits is likely due to the fact that all of these phenotypes are indirectly affected by many physiologic processes. Of these, capsule formation is the trait most likely to share physiologically direct connections to CO_2_ fitness because it is a well-characterized inducer of capsule formation in vivo (16).

Finally, we compared the set of mutants with reduced CO_2_ fitness to the set of mutants with reduced fitness in a mouse model of cryptococcosis as recently reported by Boucher et al (13). The Venn diagram in Fig. 1E shows that 137/243 mutants with reduced CO_2_ fitness in vitro also have reduced fitness during infection, further supporting the conclusion that the ability of a strain to tolerate host levels of CO_2_ is important for its ability to cause mammalian infection.

### The RIM101 and PKA pathways negatively regulate CO_2_ tolerance in an independent manner

We have previously shown that loss of both the Rim101 and PKA pathways increases CO_2_ fitness (12). The set of genes whose mutants was associated with increased CO_2_ fitness was enriched for those involved in response to alkaline pH (Fig. 1D). In addition to re-identifying the *rim101*Δ mutant, deletion mutants of components of the pathway required for proteolytic processing (Rim13 and Rim20), and hence activation, of Rim101 were also found to have increased fitness in 5% CO_2_. Activation of the Rim101 pathway has been linked to <u>E</u>ndosomal <u>S</u>orting <u>C</u>omplex <u>R</u>equired for <u>T</u>ransport (ESCRT) complex function in model yeasts (17). Furthermore, the Kronstad (18) and Alspaugh (15) labs have shown that the ESCRT complex is involved in Rim101 activation in *C. neoformans* as well. In keeping with these observations, deletion mutants of five genes with homology to ESCRT complex components (*VPS22*/23/*25*/*28*&*36*) showed increased fitness in CO_2_.

We re-tested this set of ESCRT complex mutants along with mutants of non-ESCRT complex *VPS* genes (Fig. 2A). All five ESCRT complex mutants were more fit in CO_2_ than H99 while non-ESCRT *VPS* mutants either had reduced CO_2_ fitness (*VPS1* and *VPS31*) or no phenotype (*VPS29/35/&41*). Loss of the Rim101 pathway causes reduced fitness in alkaline pH and we tested the growth of both ESCRT and non-ESCRT *VPS* mutants under these conditions. As shown in Fig. 2B, all mutants had growth defects at elevated pH in both ambient and elevated CO_2_. Therefore, it appears that the CO_2_ fitness phenotypes are not directly related to an impaired response to elevated pH responses but rather due to functions of the ESCRT/Rim101 pathway specific to CO_2_. Indeed, we have previously showed that the *rim101*Δ mutant has increased expression of a large number of genes positively regulated by the TOR pathway (12).

**Figure 2:**
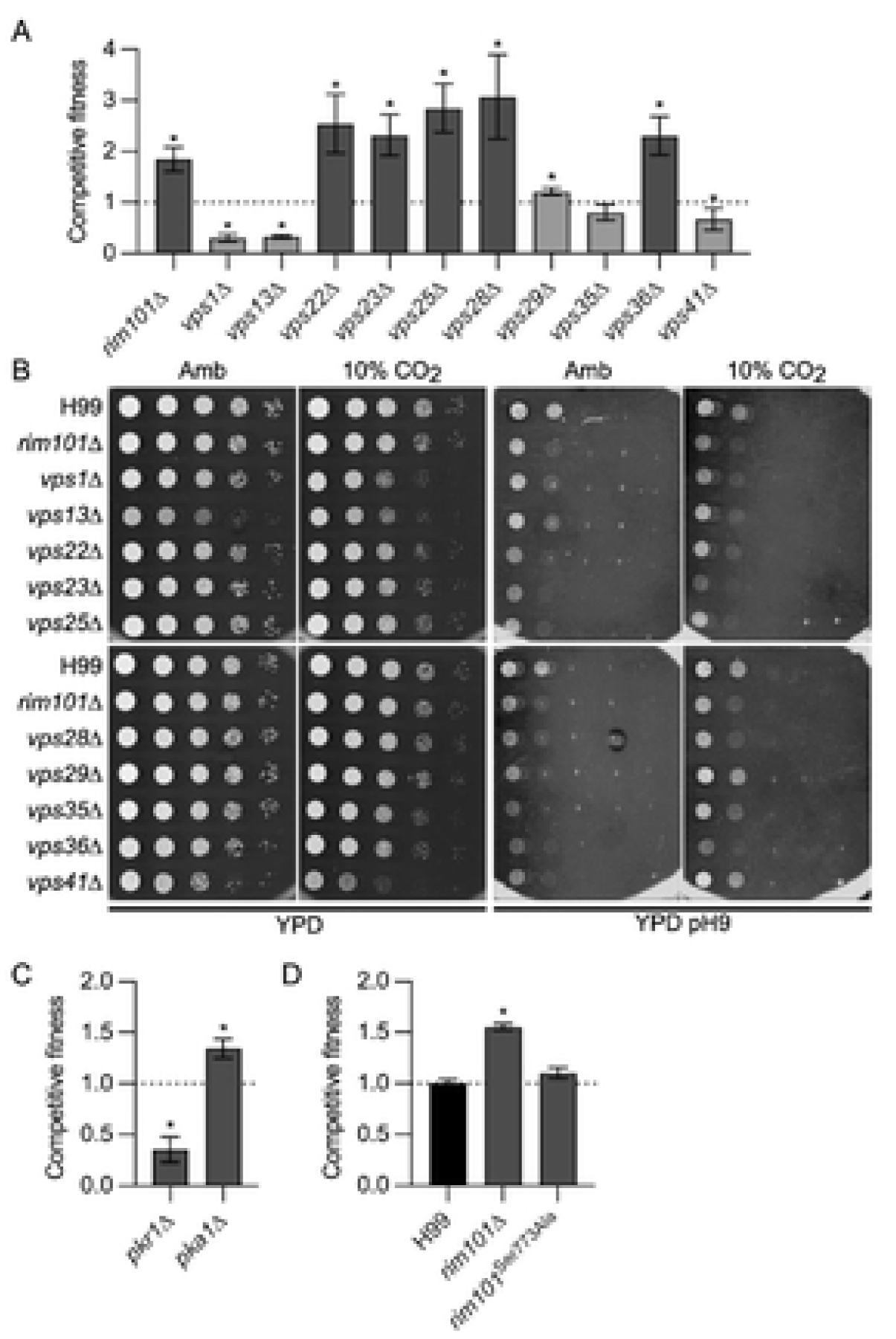
The RIM101 and PKA pathways negatively regulate CO_2_ tolerance in an independent manner. **A.** Competitive fitness was determined for the indicated vacuolar protein sorting (VPS) mutants after combining a 1:1 ratio of mNeonGreen labeled H99 and unlabeled mutant cells in RPMI 1640 medium with 165mM MOPS at pH 7 and incubating at 37°C in ambient air or 5% CO_2_ for 24 hr. Cell populations were characterized by flow cytometry and the percentage of mNeonGreen negative cells in 5% CO_2_ was divided by those in ambient conditions to calculate competitive fitness. Bars represent the average of three biological replicates with error bars indicating SD. Asterisks indicate a significant difference relative to H99 as determined by Student’s t-test (P < 0.05). **B**. Overnight cultures of *vps* and *rim101*Δ mutants were washed and concentration determined by OD_600_ reading. Strains were diluted to a starting OD_600_:1 and serial 10-fold dilutions were performed. Three µL per replicate were spotted onto standard YPD agar plates or YPD buffered with 150 mM HEPES and adjusted to pH 9. Plates were incubated at 37°C in ambient air or 10% CO_2_ for 48hr (YPD) or 144hr (YPD pH 9) before images were acquired. Images are representative of three biological replicates. Competitive fitness of *pkr1*Δ and *pka1*Δ mutants (**C**) and *rim101* mutants (**D**) in 5% CO_2_.

O’Meara et al. have demonstrated that some functions of the Rim101 pathway are regulated by the <u>P</u>rotein <u>K</u>inase <u>A</u> (PKA) pathway (19). Previously, we showed that, like loss of function mutants in the RIM101 pathway, deletion of *PKA1* increases CO_2_ fitness (12); an observation we re-confirmed in this screen (Fig. 2C). The PKA pathway is negatively regulated by Pkr1 and, consistent with that relationship, the *pkr1*Δ mutant is less fit in CO_2_ (Fig. 2C). Therefore, we tested whether the PKA pathway directly regulates Rim101 during CO_2_ stress. To do so, we took advantage of a strain constructed by O’Meara et al. in which the only source of *RIM101* is an allele in which the PKA phosphosite (Ser773) was mutated to alanine (*rim101^Ser773Ala^*, ref. 19). The *rim101^Ser773Ala^* mutant did not show a significant difference from WT in fitness under 5% CO_2_ (Fig. 2D). Therefore, both the PKA and RIM101 pathways negatively affect CO_2_ tolerance, but they do so in a manner independent of each other.

### Regulation of membrane homeostasis through both the TOR and UPR pathways is required for CO_2_ tolerance

The TOR pathway is also a key positive regulator of CO_2_ tolerance and prior work has shown that it regulates the remodeling plasma membrane asymmetry in response to elevated CO_2_ concentrations (12). Further validating the importance of the TOR pathway to CO_2_ tolerance, two putative components of the TOR complex, *SIN1* and a protein homologous to Avo2 (CNAG_02727), as well as *SIT4*, a phosphatase that appears to function downstream of TOR in *S. cerevisiae* (20), were among the mutants with reduced fitness (Table S1). A key function of the Tor-Ypk1 axis in model yeast is the regulation of sphingolipid biosynthesis (21). In *S. cerevisiae*, Ypk1 phosphorylates and activates the serine-palmityl transferase (SPT, ref. 22) which regulates the first committed step of sphingolipid biosynthesis (Fig. 3A). As previously reported, the potency of the SPT inhibitor, myriocin, against *C. neoformans* is increased in 5% CO_2_, indicating sphingolipid synthesis is required for CO_2_ tolerance (8, 12). In addition, Lee et al. have previously demonstrated that Ypk1 is required for sphingolipid biosynthesis in *C. neoformans* (23).

**Figure 3:**
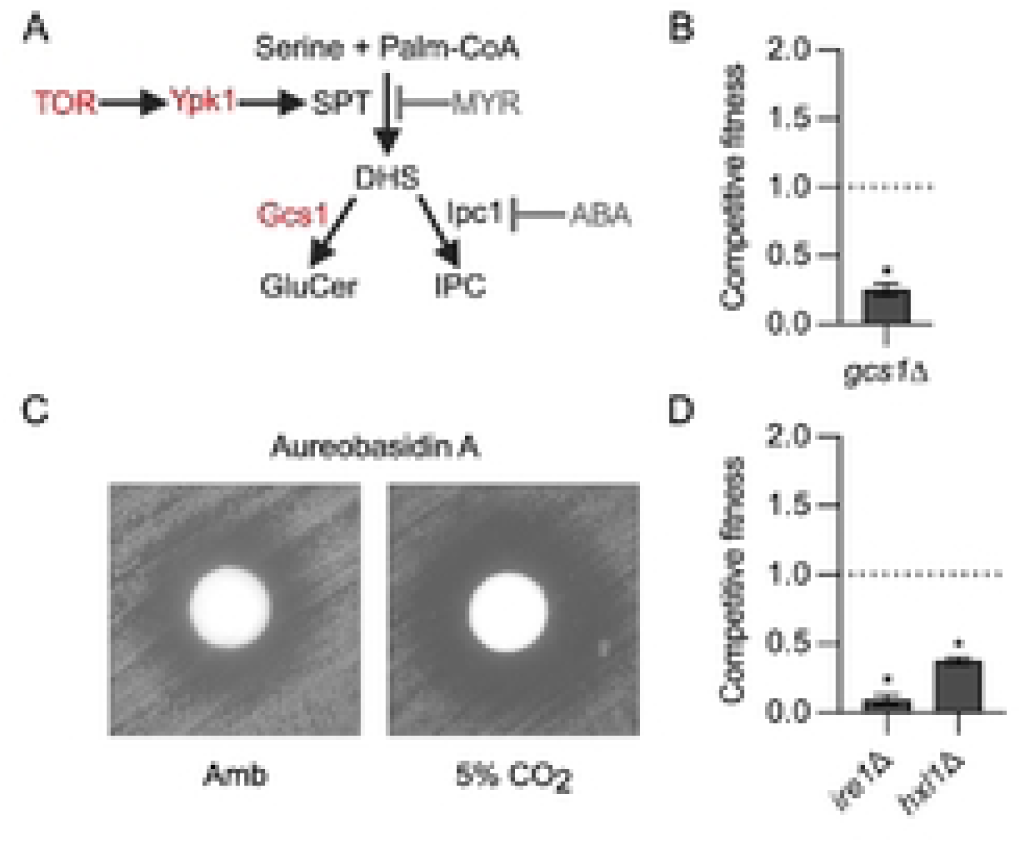
Regulation of membrane homeostasis through both the TOR and UPR pathways is required for CO_2_ tolerance. **A**. Schematic representation of the early steps in *C. neoformans* sphingolipid biosynthesis pathway. **B**. Competitive fitness was determined for glucoceramide synthase mutant *gcs1*Δ after combining a 1:1 ratio of mNeonGreen labeled H99 and unlabeled mutant cells in RPMI 1640 medium with 165mM MOPS at pH 7 and incubating at 37°C in ambient air or 5% CO_2_ for 24hr. Cell populations were characterized by flow cytometry and the percentage of mNeonGreen negative cells in 5% CO_2_ was divided by those in ambient conditions to calculate competitive fitness. Bars represent the average of three biological replicates with error bars indicating SD. Asterisks indicate a significant difference relative to H99 as determined by Student’s t-test (P < 0.05). **C**. Cells from overnight cultures of H99 were spread on solid RPMI agar plates buffered with 165mM MOPS to pH 7. Sterile disks were placed on plates and 100 µg aureobasidin A was added to each disk. Plates were incubated at 37°C in ambient air or 5% CO_2_ for 48hr before images were acquired. Images are representative of three biological replicates. **D**. Competitive fitness of the unfolded protein response (UPR) mutants *ire1*Δ and *hxl1*Δ in 5% CO_2_.

Downstream of SPT (Fig. 3A), the sphingolipid biosynthesis pathway has two branches: glucosylceramide (GC) and inositol-phosphoryl-ceramide (IPC). We identified the *gcs1*Δ mutant as having reduced CO_2_ fitness, demonstrating the importance of the GC branch in CO_2_ tolerance (Fig. 3B). IPC synthase (Ipc1) is an essential gene (24) and, therefore, to determine if the IPC branch is also required for CO_2_ tolerance, we compared the potency of the specific IPC inhibitor, aureobasidin A in ambient and 5% CO_2_. Indeed, the zone of clearance induced by aureobasidin A is increased at 5% CO_2_ relative to ambient air (Fig. 3C). These data firmly establish that an essential function of the Tor-Ypk1 axis during CO_2_ stress is to maintain sphingolipid homeostasis and that both major sphingolipid classes contribute to CO_2_ tolerance.

Membrane homeostasis is also critically dependent on the function of the endoplasmic reticulum (ER) for both the synthesis of lipids such as sterols, for the delivery to membrane proteins, and for the generation of the cell wall which buffers the cells against the effects of membrane-directed stressors. The unfolded-protein response (UPR) pathway functions to maintain ER homeostasis in the presence of a variety of stressors that affect its function (25). The two key proteins in this pathway are the kinase *IRE1* and the downstream transcription factor *HXL1* (aka *BZP1*). Both of components of the UPR pathway have reduced fitness in 5% CO_2_ (Fig. 3D). These data contribute to the growing body of evidence that host CO_2_ exerts a significant stress on the membranes of *C. neoformans* and, consequently, key mediators of membrane biosynthesis and membrane stress response pathways are critical for the ability of *C. neoformans* to replicate under those conditions.

### Host CO_2_ concentrations remodel *C. neoformans* carbon metabolism

Through its conversion to bicarbonate, CO_2_ is an essential substrate for many key reactions in central carbon metabolism (26). At the same time, it is generated as a byproduct in other reactions of central carbon metabolism such as those in the tricarboxylic acid cycle in the mitochondria. Twenty-five mutants involving central carbon metabolic processes were identified as being required for CO_2_ fitness (Table 1). The sixteen of these genes are associated with mitochondrial functions. In addition, the deletion mutants of two non-essential genes involved in glycolysis (*FBP26* and *PFK1*) have reduced fitness as do five genes associated with the synthesis of acetyl-CoA or CoASH (*YHM2*, *CAB3*, *LSC1*, *POT1*, and *ALD4*). Lastly, multiple genes in the kynurenine pathway, which generates nicotinamide, were also less fit in 5% CO_2_.

**Table 1.** Carbon metabolism genes required for fitness in host concentrations of CO_2_.

| **Role in Carbon Metabolism** | **ORF** | **Gene Name** | **Function** | **Fitness** | **Student's t-test** |
| --- | --- | --- | --- | --- | --- |
| Mitochondria | CNAG_03301 | *TIM13* | Inner membrane | 0.15 | 0.001 |
| Mitochondria | CNAG_02974 | *POR2* | Porin | 0.38 | 0.011 |
| Mitochondria | CNAG_05573 | *COX17* | Cytochrome Oxidase | 0.49 | 0.016 |
| Mitochondria | CNAG_03298 | *TIM11* | Inner membrane | 0.49 | <0.0001 |
| Mitochondria | CNAG_05267 | *none* | NADH Dehydrogenase subunit | 0.55 | 0.014 |
| Mitochondria | CNAG_05395 | *VAM6* | GEF | 0.55 | 0.02 |
| Mitochondria | CNAG_02575 | *YPT7* | GTPase | 0.56 | 0.032 |
| Mitochondria | CNAG_03375 | *LSC1* | Succinyl-CoA Synthetase | 0.56 | 0.025 |
| Mitochondria | CNAG_02833 | *none* | NADH Dehydrogenase subunit | 0.59 | 0.042 |
| Mitochondria | CNAG_06663 | *QCR6* | Ubiquinol-Cytochrome C reductase | 0.60 | <0.0001 |
| Mitochondria | CNAG_02905 | *COA6* | Cytochrome C oxidase assembly | 0.64 | 0.013 |
| Mitochondria | CNAG_02403 | *none* | NADH Dehydrogenase Ubiquinone subcomplex 9 | 0.70 | 0.043 |
| Mitochondria | CNAG_00440 | *SSN801* | Cyclin-like subunit | 0.70 | 0.014 |
| Mitochondria | CNAG_07373 | *URA2* | Carbamoyl phosphate synthase | 0.60 | 0.012 |
| Mitochondria | CNAG_03269 | *ALD4* | Aldehyde dehydrogenase | 0.47 | 0.014 |
| Glycolysis | CNAG_04221 | *FBP26* | Fructose 2,6-bisphosphatase | 0.41 | 0.01 |
| Glycolysis | CNAG_04676 | *PFK1* | 6-phosphofructokinase | 0.52 | 0.027 |
| Nicotinamide Biosynthesis | CNAG_02793 | *BNA4* | Kynurenine 3-mono-oxygenase | 0.15 | 0.003 |
| Nicotinamide Biosynthesis | CNAG_02868 | *BNA2* | tryptophan 2,3-dioxygenase | 0.15 | 0.001 |
| Nicotinamide Biosynthesis | CNAG_02780 | *BNA5* | Kynureninase | 0.19 | 0.002 |
| Nicotinamide Biosynthesis | CNAG_02779 | *BNA6* | Nicotinate-nucleotide diphosphorylase | 0.27 | 0.002 |
| Acetyl-CoA Biosynthesis | CNAG_00490 | *POT1* | Acetyl-CoA acetyl transferase, β-oxidation | 0.62 | 0.036 |
| Acetyl-CoA Biosynthesis | CNAG_07348 | *CAB3* | Phosphopantothenoyl-cysteine decarboxylase | 0.40 | 0.004 |
| Acetyl-CoA Biosynthesis | CNAG_06096 | *YHM2* | Citrate/α-ketoglutarate anti-porter | 0.19 | 0.005 |
| Acetyl-CoA Biosynthesis | CNAG_04531 | *none* | Enoyl CoA hydratase, β-oxidation | 0.60 | 0.017 |
| Acetyl-CoA Biosynthesis | CNAG_05721 | *MFE2* | Multi-functional protein involved in β-oxidation | 0.69 | 0.0269 |

To further explore the effect of host CO_2_ concentrations on the metabolic state of *C. neoformans*, we compared the non-lipid metabolomic profile of *C. neoformans* incubated in buffered RPMI at ambient CO_2_ to that from cells incubated in 5% CO_2_ after 24hr at 37°C. The abundance of 90 molecules differed between 5% CO_2_ and ambient air (± 1 log_2_, adjust p <0.05, Table S2). Strikingly, glucose; two glycolysis intermediates, phosphoenopyruvate (PEP) and dihydroxyacetone-3-phosphate (DHAP); and fructose (Fig. 4A) are reduced, suggesting that exposure to 5% CO_2_ depletes glucose and may lead to decreased flux through glycolysis. 5% CO_2_ is a strong inducer of the polysaccharide capsule in *C. neoformans* (16) and we have previously shown that genes involved in capsule biosynthesis are enriched in the set of genes upregulated by exposure of H99 to host levels of CO_2_ (12). The carbohydrates that make up the capsule are derived from glucose, suggesting that CO_2_ conditions divert glucose from glycolysis to capsule biosynthesis (27). Indeed, the levels of a glucose-derived capsule biosynthesis intermediate, UDP-glucuronate (UDP-GlnA), are reduced 4-fold in 5% CO_2_ relative to ambient CO_2_ levels (Fig. 4B), supporting the idea that flux through this pathway is increased. In the next step of capsule biosynthesis, UDP-GlnA is converted to UDP-xylose by UDP-xylose synthase (*UXS1*, ref. 28). Consistent with this function the *uxs1*Δ mutant has reduced capsule thickness (29). Deletion of *UXS1* increases CO_2_ fitness (Fig. 4C), suggesting that interrupting capsule biosynthesis early in the pathway provides a growth advantage. In contrast, deletion of *UGE1*, the enzyme that epimerizes UPD-glucose to UDP-galactose, increases capsule thickness (30) and reduces CO_2_ fitness (Fig. 4C). Taken together, the induction of capsule biosynthesis may alter glucose availability and loss of genes involved in the early steps of capsule biosynthesis may indirectly affect CO_2_ tolerance by altering glucose flux through that pathway.

**Figure 4:**
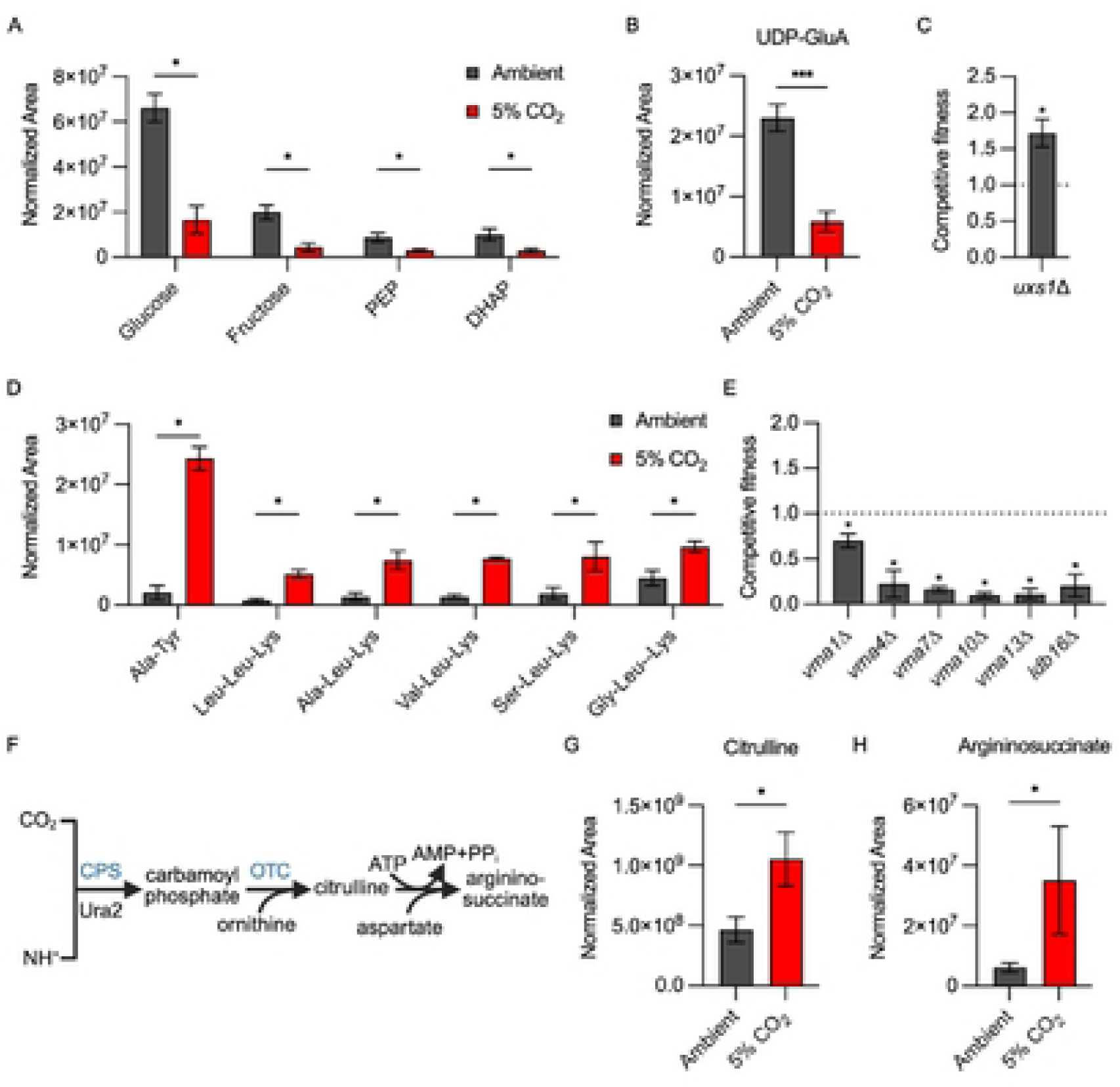
Host CO_2_ concentrations remodel *C. neoformans* carbon metabolism. The normalized area of each metabolite was determined by tandem mass spectrometry as described in materials and methods for H99 incubated in buffered RPMI in ambient air or 5% CO_2_. Asterisks indicate statistically significant differences in abundance at 5% CO_2_ relative to ambient air (Welch corrected t test, p <0.05). **A**. Relative amounts of glycolytic intermediates. **B**. Relative amounts of UDP-Glucuronic Acid (UDP-GluA). **C**. Competitive fitness of the *uxs1*Δ and *uge1*Δ mutants relative to H99 was determined for indicated strains after combining a 1:1 ratio of mNeonGreen labeled H99 and unlabeled mutant cells in RPMI 1640 medium with 165mM MOPS at pH 7 and incubating at 37°C in ambient air or 5% CO_2_ for 24 hr. Cell populations were characterized by flow cytometry and the percentage of mNeonGreen negative cells in 5% CO_2_ was divided by those in ambient conditions to calculate competitive fitness. Bars represent the average of three biological replicates with error bars indicating SD. Asterisks indicate a significant difference relative to H99 as determined by Welch corrected, two-tailed t-test (P < 0.05). **D**. Relative abundance of the indicated di- and tripeptides in ambient and 5% CO_2_. **E**. Competitive fitness of mutants involved in vacuolar ATPase function. **F**. Schematic of the initial steps of the urea cycle in *C. neoformans*.

Metabolomic studies in other yeast have shown that depletion of glucose increases protein catabolism in order to mobilize amino acids as carbon sources (31). As shown in Fig. 4D, nine di- and tripeptides are increased in 5% CO_2_, a strong indication that *C. neoformans* catabolizes proteins under these conditions. In addition, the amounts of six amino acids (Leu, Phe, Tyr, Trp, His, and Met) are increased by ≥ 2-fold in cells grown in 5% CO_2_ (Table S2). Of the nine di- and tri-peptides identified, all contain ketogenic amino acids and 7/9 peptides contain Leu and Lys which are exclusively ketogenic amino acids. Ketogenic amino acids can be further catabolized in the mitochondria to acetyl-CoA which then enters the TCA cycle as an alternative to acetyl-CoA derived from glycolysis-derived pyruvate (32). Thus, it appears that *C. neoformans* mobilizes alternative carbon nutrients by protein catabolism to compensate for reduced amounts of glucose after shift to host levels of CO_2_.

Protein catabolism occurs in the vacuole and is dependent upon vacuolar ATPase (V-ATPase) to generate the acidic environment necessary for the activation of vacuolar peptidases (33). Multiple strains lacking genes coding for V-ATPase components or proteins involved in the assembly of the V-ATPase complex show reduced CO_2_ fitness (Fig. 4E), indicating that maintenance of vacuolar pH is critical for tolerance of high CO_2_. Although V-ATPase mutants have pleomorphic effects on the cell, protein catabolism is clearly occurring during the initial phases of CO_2_ stress and loss of V-ATPase function would likely interfere with efficient generation of ketogenic amino acids necessary to maintain mitochondrial respiration.

CO_2_/HCO_3_ ^-^ is essential for the cell, but high levels of CO_2_ are stressful and can ultimately be toxic. As such, CO_2_ is both generated and consumed by metabolic reactions in the cell. The urea cycle is one of the CO_2_-consuming processes utilized by eukaryotic cells (Fig 4F). Ura2 is a carbamoyl phosphate synthase that catalyzes the rate-limiting condensation of CO_2_ with ammonia which feeds into the urea cycle. Previous work by our group had identified that deletion of *URA2* reduces CO_2_ fitness and our screen confirmed this result (Table 1). Carbamoyl-phosphate enters the urea cycle by reacting with ornithine to generate citrulline which, in turn, is converted to argininosuccinic acid (Fig. 4F). Both the citrulline and argininosuccinic acid levels are elevated at 5% CO_2_ relative to ambient levels (Fig. 4G&H), suggesting increased conversion of CO_2_ to urea cycle intermediates relative to ambient CO_2_. Supporting this conclusion, in *S. cerevisiae*, urea cycle activity is enhanced under increased CO_2_ concentrations to deplete excess CO_2_/HCO_3_ (34). Ura2 is also required for the synthesis of uridine/uracil and deletion mutants are auxotrophic for those nutrients. However, Chadwick et al. found that the reduced fitness of the *ura2*Δ mutant in 5% CO_2_ was not rescued by uracil or uridine supplementation while the auxotrophic phenotypes were reversed (11). Therefore, Ura2 may buffer the cell against high levels of CO_2_ by converting it into intermediates of the urea cycle. Together, these data indicate that host levels of CO_2_ lead to substantial alteration of central carbon metabolism through multiple mechanisms.

### Host concentrations of CO_2_ alter *C. neoformans* cellular redox balance

In *S. cerevisiae*, acute depletion of glucose leads to an increase in mitochondrial respiration. Based on these precedents and the large number of mitochondrial genes required for CO_2_ fitness, we predicted that host CO_2_ concentrations would increase mitochondrial respiration and membrane potential. To test this hypothesis, we characterized the membrane potential of mitochondria in ambient and 5% CO_2_ using JC-1 staining as previously reported by the Kronstad lab (35). In this assay, increased mitochondrial membrane potential leads to a red shift in the JC-1 dye. As shown in Fig. 5A, growth in 5% CO_2_ leads to an increase in the proportion of red-shifted cells relative to ambient CO_2_. This increase in mitochondrial membrane is consistent with an increase in respiration at host concentrations of CO_2_ (36). The membrane potential is driven by the TCA cycle and electron transport chain enzymes. Accordingly, deletion mutants of the TCA cycle enzyme succinyl CoA synthetase (*LSC1*) as well as three complex I/ubiquinone NADH dehydrogenase components CNAG_05267, CNAG_02403, and CNAG_02833 have reduced fitness in host CO_2_ concentrations (Table 1). *PFK1* and *FBP26* reduce the ability of yeast to transition from glycolysis to respiration (37). We, therefore, suggest this function may contribute to the reduced fitness of the *pfk1*Δ and *fbp26*Δ mutants (Table 1) in 5% CO_2_.

**Figure 5:**
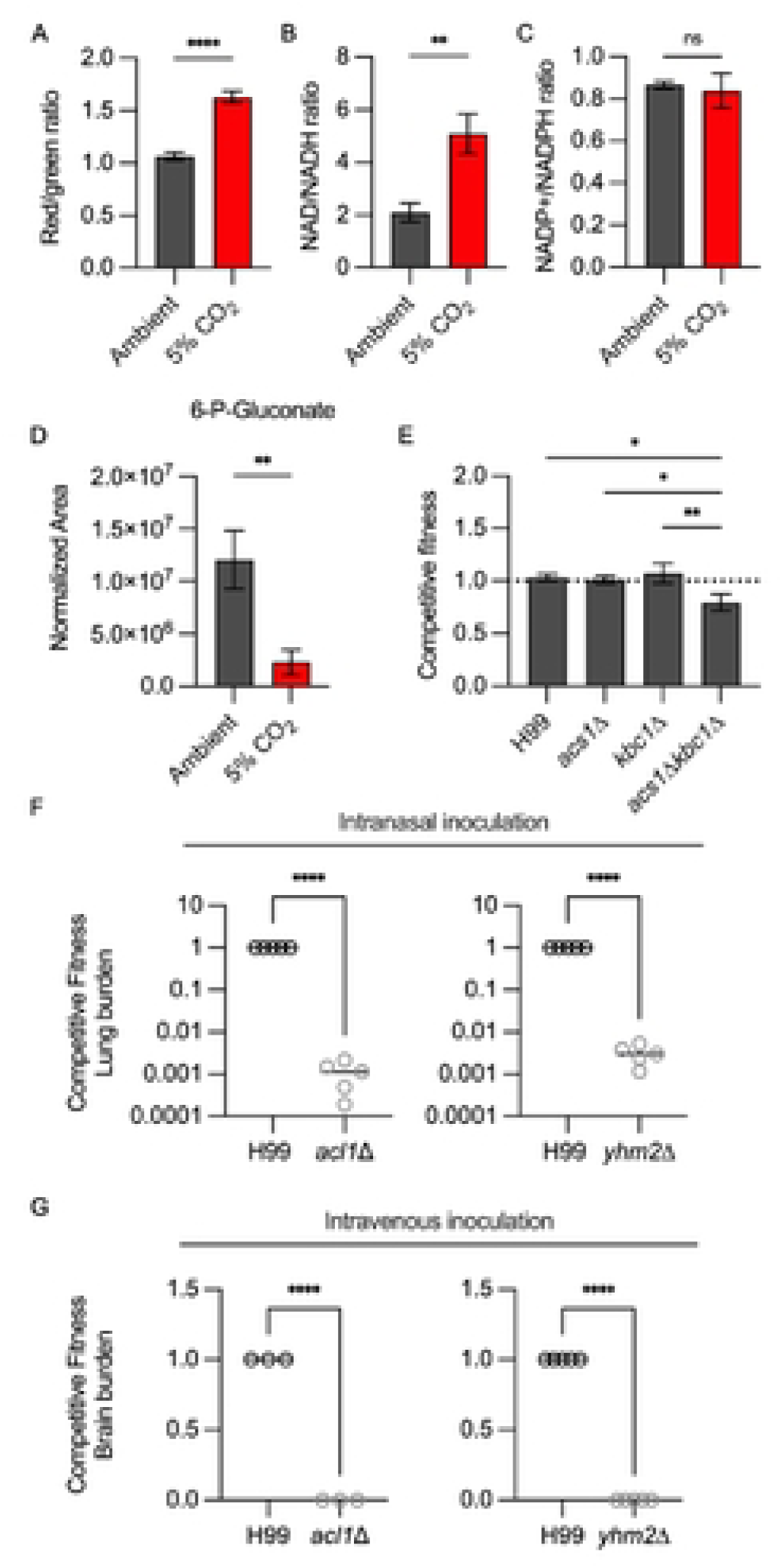
Host concentrations of CO_2_ alter *C. neoformans* cellular redox balance and engage multiple acetyl-CoA generating pathways for tolerance. **A.** The mitochondrial membrane potential of H99 in ambient or 5% CO_2_ conditions for 24hr was assessed by JC-1 staining. JC-1 staining was analyzed by dividing the MFI resulting from excitation with the blue (488nm) laser in the BL2 emission filter (574/26) by the MFI in the BL1 emission filter (530/30), i.e. red/green ratio of single yeast cells. Bars represent the average red/green ratio of a minimum of 10,000 cells per condition and error bars represent the SEM. Asterisks indicate a significant difference relative to H99 as determined by Student’s t-test (P < 0.05). Cellular stores of NAD/NADH (**B**) and NADP+/NADPH (**C**) were determined for H99 after incubation in buffered RPMI medium at ambient or 5% CO_2_ conditions for 24hr. Data are presented as ratios of the oxidized or reduced forms. Bars represent the average of three biological replicates and SD with significance relative to ambient conditions as determined by unpaired t-test (P < 0.05). D) The metabolic profile of H99 grown for 24hr in ambient air or 5% CO_2_ was analyzed by tandem mass spectrometry. Data were processed and are presented for 6-P-Gluconate as normalized area. Asterisk indicates p <0.05 for the comparison of ambient to 5% CO_2_ using two-tailed Welch corrected t test. **E**. Competitive fitness was determined for indicated strains after combining a 1:1 ratio of mNeonGreen labeled H99 and unlabeled mutant cells in RPMI 1640 medium with 165mM MOPS at pH 7 and incubating at 37°C in ambient air or 5% CO_2_ for 24 hr. Cell populations were characterized by flow cytometry and the percentage of mNeonGreen negative cells in 5% CO_2_ was divided by those in ambient conditions to calculate competitive fitness. Bars represent the average of three biological replicates with error bars indicating SD. Asterisks indicate a significant difference relative to H99 as determined by one-way ANOVA with Tukey’s multiple comparisons test (P < 0.05). The competitive fitness of H99 and indicated strains (*acl1*Δ or *yhm2*Δ) was determined in vivo after two inoculation routes in CD-1 mice. **F**. For intranasal inoculation (IN), wild-type and mutant cells were mixed in a 1:1 ratio prior to inoculation with 5 × 10^4^ CFU/animal in a 50 µL volume. **G**. For intravenous inoculation (IV), wild-type and mutant cells were mixed in a 1:1 ratio prior to inoculation with 1 × 10^5^ CFU/animal in 100 µL volume. Organs were harvested at day 4 post-inoculation. Tissues were homogenized and serially diluted on YNB or YNB with 400 µg/mL hygromycin and incubated at 30°C for two days before counting CFUs. The competitive fitness was determined by dividing the CFU of mutant cells (HYG resistant) by the CFU of H99 cells in lung (F) from the IN model or brain (G) from the IV model. Data are represented with the average CFU/mouse and SD. Asterisks indicate a significant difference relative to H99 as determined by unpaired t-test (P < 0.05).

To further characterize the effect of CO_2_ on the redox state of *C. neoformans*, we measured the ratios of NAD+/NADH and NADP+/NADPH under ambient and host CO_2_ concentrations. As shown in Fig. 5B, CO_2_ increases the NAD+/NADH ratio by ∼2.5 fold. The increased NAD+/NADH ratio is consistent with the higher mitochondrial membrane potential observed in Fig. 5A and further supports the conclusion that 5% CO_2_ increases respiration and TCA cycle activity relative to ambient CO_2_ concentrations. One of the consequences of increased respiration is increased oxidative stress in the cell. NADPH is a critical component of the biochemical processes that detoxify ROS (38). Interestingly, the ratio of NADP+/NADPH is unchanged at 5% CO_2_ compared to ambient CO_2_ (Fig. 5C). This suggests that CO_2_-tolerant *C. neoformans* can compensate for the increase in respiration and oxidative state of the cell at 5% CO_2_ without depleting NADPH stores significantly. The pentose-phosphate pathway generates two molecules of NADPH through the oxidation of glucose first to 6-phospho-gluconate and then, through an oxidative decarboxylation step, to ribulose-5-phosphate. The levels of 6-phosphogluconate are reduced in 5% CO_2_ relative to ambient CO_2_ conditions which is consistent with increased pentose phosphate pathway activity (Fig. 5D), possibly to maintain NADPH levels and compensate for increased ROS resulting from increased mitochondrial respiration.

### Multiple acetyl-CoA generating pathways are required for *C. neoformans* tolerance of host CO_2_ concentrations

Acetyl-CoA is not only required for mitochondrial respiration through the TCA cycle but is also a fundamental substrate for lipid biosynthesis as well as for histone acetyl transferases involved in the regulation of gene expression (39). Acetyl-CoA cannot diffuse across organelle membranes and, therefore, distinct pools of acetyl-CoA are present in the mitochondria, cytosol and nucleus. In the presence of glucose, glycolysis delivers pyruvate-derived acetyl-CoA to the mitochondria and TCA cycle. Our metabolomics data suggest that glucose and glycolysis are reduced in 5% CO_2_ and that ketogenic amino acids are mobilized to provide acetyl-CoA to support the increased mitochondrial respiration driven by the acute reduction in glucose.

Lipid catabolism through β-oxidation also contributes to the acetyl-CoA pool and, therefore, could also contribute to acetyl-CoA homeostasis during CO_2_ stress (31). Supporting this hypothesis, we identified three mutants in the β-oxidation pathway that have reduced CO_2_ fitness: the enoyl CoA hydratase CNAG_04531, the multi-functional protein *MFE2*, and the 3-oxo-acyl CoA thiolase *POT1* (Table 1). Enoyl-CoA hydratases catalyze the second step of fatty acid β-oxidation while Pot1 directly generates acetyl-CoA. *MFE2* is a peroxisomal enzyme that catalyzes the first two steps of β-oxidation (38). *MFE2* is subject to glucose repression in *C. neoformans* and *POT1* has been shown to be negatively regulated by glucose in *S. cerevisiae* (41). The expression of both *MFE2* (1.8-fold, FDR = 9e-15) and *POT1* is (3-fold, FDR 1e-68) is increased in 5% CO_2_, further supporting its likely role in generating acetyl-CoA under these conditions (12). In addition, β-oxidation is positively regulated by the AMP-activated protein kinase Snf1 in other systems (42) and this kinase is required for full fitness at host CO_2_ concentrations (competitive index: 0.67; p = 0.012, Table S1). Finally, *POT1*, *MFE2*, and enoyl CoA hydratase CNAG_04531 are required for full fitness during infection (13). Thus, the in vivo fitness defects of genes related to β-oxidation may be due, at least in part, to their importance in the adaptation of *C. neoformans* to host CO_2_.

Two additional enzymes could also contribute to the generation of acetyl-CoA: acetyl-CoA synthetase (*ACS1*) and aceto-acetyl-CoA synthetase (*KBC1*, ref. 43). Acs1 converts acetate to acetyl-CoA. Kbc1 condenses the ketone body acetoacetate with CoA to give aceto-acetyl-CoA which, in turn, is a substrate for Pot1, leading to two molecules of acetyl-CoA. We, therefore, tested the CO_2_ fitness of *acs1*Δ, *kbc1*Δ, and the corresponding double deletion mutants using the competition assay. The *kbc1*Δ *acs1*Δ double mutant showed a modest reduction in CO_2_ fitness while neither of the single mutants differed from WT (Fig. 5E), suggesting that these enzymes play a minor role in maintaining acetyl-CoA homeostasis during CO_2_ stress.

Amino acid catabolism and β-oxidation to acetyl-CoA occur in the mitochondria and directly provide acetyl-CoA for use in the TCA cycle. However, acetyl-CoA is also required in other cellular components to support biosynthesis and gene expression. An important source of cytosolic and nuclear acetyl-CoA in *C. neoformans* and other cells is, in fact, the mitochondria. Indeed, previous work by the Kronstad lab showed that the majority of cytosolic and nuclear acetyl-CoA appears to be derived from citrate generated in the mitochondria (44). Citrate is exported to the cytosol, where it is converted to acetyl-CoA and oxaloacetate by ATP-citrate lyase (*ACL1*). In prior work, we found that the *acl1*Δ mutant has a severe growth defect in 5% CO_2_ (43). This suggests that nuclear and cytosolic acetyl-CoA stores remain primarily dependent on Acl1.

The deletion mutant of a tricarboxylic acid transporter, CNAG_06096, has a strong fitness defect in 5% CO_2_ (Table 1). CNAG_06096 is homologous to *S. cerevisiae* Yhm2 which is a mitochondrial citrate-oxoglutarate antiporter (45). In other fungi with ATP-citrate lyase enzymes such as *Aspergillus* spp., deletion of the citrate-oxoglutarate transporter leads to reduced acetyl-CoA production (46). Presumably, this is due to reduced cytosolic citrate in the absence of the transporter. If Yhm2 plays this role during mammalian *C. neoformans* infection, then, like the *acl1*Δ mutant, it should have a profound fitness defect in vivo. To test this hypothesis, we infected mice with a 1:1 mixture of H99 and either the *acl1*Δ or *yhm2*Δ mutant via inhalation (Fig. 5F) and intravenous injection (Fig. 5G). As expected, based on the previous report of its reduced virulence, the *acl1*Δ had dramatically reduced fitness in the lung and the brain. The *yhm2*Δ mutant showed a similarly profound reduction in fitness relative to WT. These data indicate that mitochondrial citrate-derived acetyl-CoA is required for *C. neoformans* tolerance of CO_2_ and infection of mammals.

Taken together, these data indicate that host concentrations of CO_2_ establish a relatively low glucose state in *C. neoformans*. In response, oxidation of fatty acids and amino acids generates mitochondrial acetyl-CoA which then supports increased respiratory metabolism. Cytosolic and nuclear pools of acetyl-CoA remain largely dependent on the export of mitochondrial citrate through the transporter Yhm2; citrate is then converted to acetyl-CoA by Acl1 in the nucleus and cytosol.

### The glutathione and TOR-Ypk1 pathways buffer *C. neoformans* against CO_2_-induced ROS accumulation

One of the consequences of increased mitochondrial respiration is the generation of reactive oxygen species (ROS). Glutathione is an important mechanism by which ROS species are detoxified. Recently, Black et al. described the importance of glutathione-mediated redox regulation in the virulence of *C. neoformans* through effects on melanization and titan cell formation (47). In addition, loss of glutathione affects a variety of metabolic pathways and reduces fitness in low nutrient growth conditions. The glutathione synthetases *GSH1* and *GSH2* are both required for CO_2_ tolerance in H99 (Fig. 6A). The biosynthesis of glutathione is integrated with sulfate assimilation as well as methionine and cysteine biosynthesis (Fig. 6B, ref. 48). Consistent with this integration, five genes in the sulfur assimilation pathway (*SUL1*, *MET3*, *MET5*, *MET10*) are required for CO_2_ tolerance along with *CYS4* which converts homocysteine to cystathionine and *MET2* which generates the homoserine that is coupled with cystathionine to generate cysteine (Fig. 6A&B). Homoserine levels are significantly reduced under 5% CO_2_ (Fig. 6C), further supporting the notion that this pathway is highly active under those conditions.

**Figure 6:**
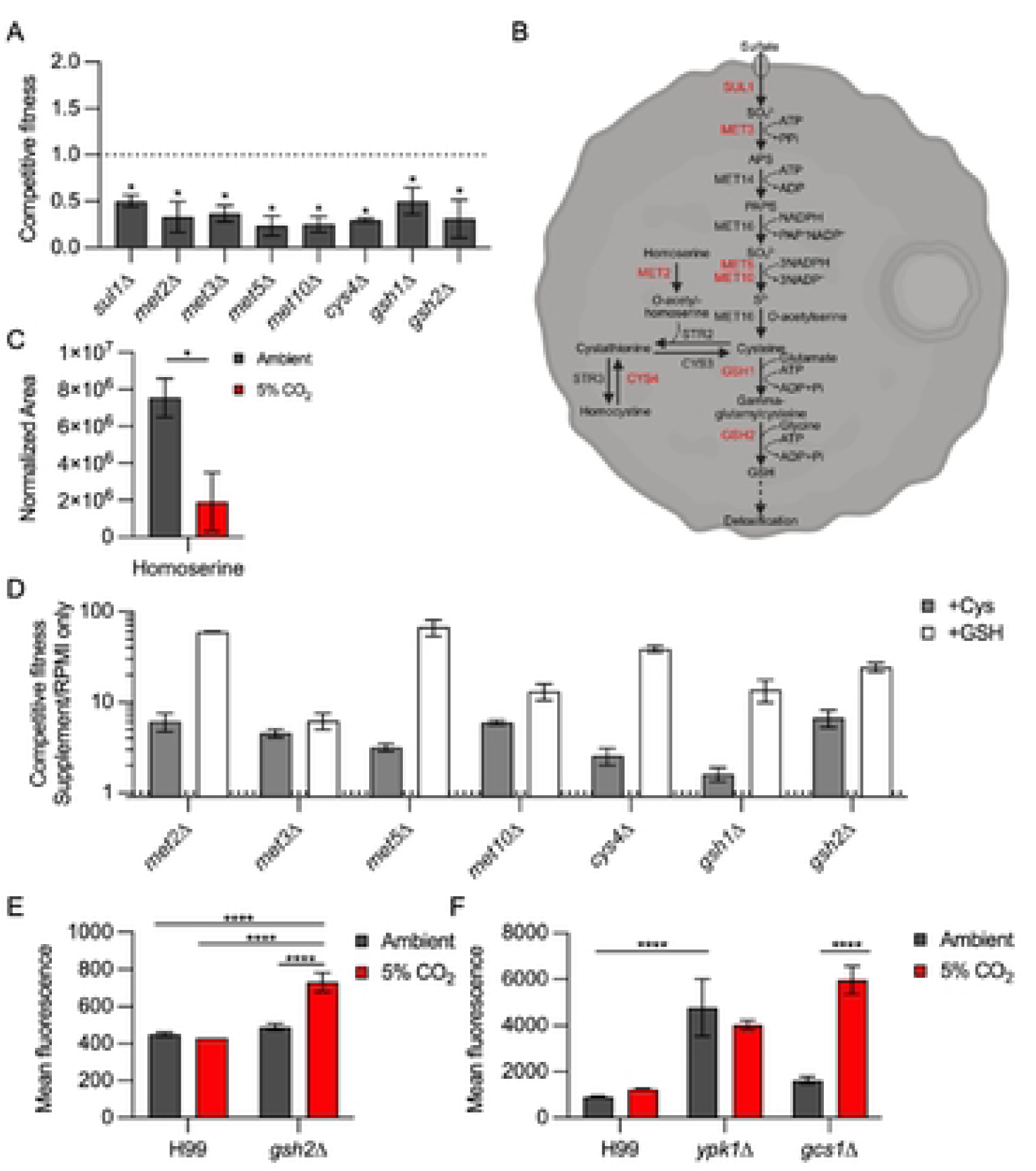
The glutathione and TOR-Ypk1 pathways buffer *C. neoformans* against CO_2_-induced ROS accumulation. **A**. Competitive fitness was determined for indicated strains after combining a 1:1 ratio of mNeonGreen labeled H99 and unlabeled mutant cells in RPMI 1640 medium with 165mM MOPS at pH 7 and incubating at 37°C in ambient air or 5% CO_2_ for 24 hr. Cell populations were characterized by flow cytometry and the percentage of mNeonGreen negative cells in 5% CO_2_ was divided by those in ambient conditions to calculate competitive fitness. Bars represent the average of three biological replicates with error bars indicating SD. Asterisks indicate a significant difference relative to H99 as determined by Student’s t-test (P < 0.05). **B**. Schematic of the glutathione biosynthesis pathway in *C. neoformans*. Genes required for CO_2_ fitness are indicated in red. **C**. The metabolic profile of H99 grown for 24 hr in ambient air or 5% CO_2_ was analyzed by tandem mass spectrometry. Data were processed and are presented for homoserine as normalized area. The normalized area of each metabolite was determined by tandem mass spectrometry as described in materials and methods for H99 incubated in buffered RPMI in ambient air or 5% CO_2_. Asterisks indicate statistically significant differences in abundance at 5% CO_2_ relative to ambient air (Welch corrected two-tailed t test, p <0.05). **D**. Competitive fitness of the glutathione biosynthesis pathway mutants was determined as in (**A**) with the following modification: cells were grown in RPMI 1640 medium with 165mM MOPS at pH 7 alone or with the addition of 200 µM cysteine or 1 mM glutathione for 24 hr in 5% CO_2_. The percentage of mNeonGreen negative cells in media with supplement was divided by the percentage of mNeonGreen negative cells without supplement to determine competitive fitness. Bars indicate the average of three biological replicate with error bars indicating SD. Asterisks indicate a significant difference relative to H99 as determined by Student’s t-test (P < 0.05). **E**&**F**. ROS activity of indicated strains after growth in ambient air or 5% CO_2_ for 24hr was assessed by DCFDA fluorescence assay. The average MFI of three biological replicates for each strain and condition is represented with error bars indicating SD. Asterisks indicate a significant difference as determined by one-way ANOVA with Sidak’s multiple comparisons test (P < 0.05).

To explore the role of these pathways in CO_2_ tolerance, we compared the CO_2_ fitness of each mutant in the presence and absence of supplemental cysteine and glutathione (Fig. 6D). The CO_2_ fitness of each mutant was improved in the presence of both cysteine and glutathione. Black et al. had found that reduced glutathione had effects on the cell beyond its role in detoxifying ROS (47). We, therefore, used the DCFDA assay to characterize the levels of ROS induced by CO_2_ in the WT strain and the *gsh2*Δ mutant (Fig 6E). In WT cells, 5% CO_2_ does not induce a significant increase in ROS. However, cellular ROS increases significantly in the *gsh2*Δ mutant relative to WT in 5% CO_2_ (Fig. 6E). These data suggest that host concentrations of CO_2_ cause oxidative stress in *C. neoformans* but that CO_2_ tolerant strains such as H99 utilize glutathione to detoxify these species as part of a compensatory response. It is also possible that glutathione is required to maintain metabolic homeostasis as well. Taken together, these data indicate that the reduced in vivo fitness of genes involved in glutathione homeostasis may also be due to their role in the response to increased CO_2_ concentrations encountered by *C. neoformans* during mammalian infection.

In *S. cerevisiae*, the TOR-Ypk1 pathway also buffers the cell from ROS generated in the mitochondria and vacuole through its regulation of sphingolipid biosynthesis (49). In *C. neoformans*, TOR-Ypk1 is required for sphingolipid biosynthesis and we hypothesized that this pathway may also buffer the cell against CO_2_-induced ROS. High levels of ROS in ambient CO_2_ are present in the *ypk1*Δ mutant relative to WT (Fig. 6F). This observation is consistent with the model proposed by Niles et al. in which TOR-Ypk1 suppresses the accumulation of ROS in *S. cerevisiae* (49). The levels of ROS in the *ypk1*Δ mutant do not increase further in 5% CO_2_. However, the loss of ROS homeostasis in the *ypk1*Δ mutant seems likely to contribute to its susceptibility to 5% CO_2_. As discussed above Ypk1 is required for synthesis of sphingolipids in *C. neoformans* and other fungi through regulation of SPT (22, 23). Niles et al. also found that reduced sphingolipid biosynthesis increased cellular ROS (49). If this mechanism is also operative at host levels of CO_2_ in *C. neoformans*, then the *gcs1*Δ mutant would be expected to have the same phenotype. Although loss of *GCS1* does not alter ROS levels in ambient CO_2_ concentrations (Fig. 6F), ROS levels are increased in the *gcs1*Δ mutant in 5% CO_2_. Thus, the TOR-Ypk1 pathway is required for the prevention of toxic levels of cellular ROS and its regulation of sphingolipid biosynthesis is likely to contribute to this function. Furthermore, the glucosylceramide arm of the sphingolipid pathway directly contributes to the buffering of ROS generated by *C. neoformans* growth at host concentrations of CO_2_.

## Discussion

To cause disease in humans, *C. neoformans* must transition from the low CO_2_, external environment to the host environment where CO_2_ concentrations are ∼100-fold higher (9). In general, strains that are unable to grow at host CO_2_ concentrations cannot cause disease in mammalian models of cryptococcosis (8, 10). To better understand the determinants of host CO_2_ tolerance, we performed a large-scale functional genomics screen to identify genes and cellular processes required for a CO_2_ tolerant strain to make the transition from CO_2_ concentrations found in the external environment to those present in the host. The set of genes which modulate CO_2_ tolerance is relatively large and encompasses a variety of cellular functions. This observation leads to the first two conclusions of this work. Specifically, the transition of *C. neoformans* to host concentrations represents a profound stress and there is no single CO_2_ stress response pathway. In previous work (12), we found that stress response pathways could be adaptive or maladaptive in the setting of host concentrations of CO_2_. Indeed, critical regulatory pathways such as the cAMP-PKA pathway, the RIM101 pathway, and the cell wall integrity MAPK pathways are maladaptive (12). In contrast, the TOR-Ypk1, calcineurin, and RAM pathways are essential during the transition from low CO_2_ to high CO_2_ concentrations (12, 50). From this near-genome wide screen, the UPR and glutathione biosynthesis pathways have been added to the list of those required for CO_2_ tolerance.

Initially, it seems almost counter-intuitive that major signaling pathways required for *C. neoformans* virulence are maladaptive under CO_2_ stress. To address this seeming conundrum, we propose that these observations are the result of a balance that is likely to occur as a tolerant cell adapts to high CO_2_. During this transition, the relative activities of the maladaptive are reduced while the activities of the adaptive pathways are increased. During infection, CO_2_ is but one stress to which the cell must adapt in order to successfully establish infection: others include pH, direct oxidative stress, nutrient stress, and temperature stress. Indeed, during temperature stress, the cell wall integrity pathway is activated while the Sch9 arm of the TOR pathway is maladaptive (51). Accordingly, the response to high CO_2_ stress is integrated into a cellular response involving multiple stressors during successful infection.

One of the main goals of this work was to move beyond regulatory pathways to cellular and physiological processes and functions that are key to the multifactorial response to host concentrations of CO_2_. With those goals in mind, we focused on two sets of genes that emerged from our screen: metabolism and ROS-related genes. With respect to the first set of genes, we integrated our genetic data with metabolomic profiling as well as assessments of mitochondrial function and redox status. From this analysis, we propose the following model for the metabolic remodeling of *C. neoformans* metabolism during the transition to host concentrations of CO_2_.

First, we suggest that the reduced steady state concentrations of glucose and glycolytic pathway intermediates is due, in part, to diversion of glucose to UDP-glucose which, in turn, is a key substrate in the synthesis of capsule. Host concentrations of CO_2_ were one of the first well-described in vitro inducers of capsule formation (16). Because capsule is an extracellular glucose-derived polymer (27), its synthesis re-routes massive amounts of metabolic resources to the extracellular space. Second, the reduction in glucose results in an increased dependence on mitochondrial respiration to support central carbon metabolism. Third, alternative carbon sources such as ketogenic amino acids and fatty acids must be catabolized to acetyl-CoA to provide sufficient carbon flux through the TCA cycle of the mitochondria. Mobilization of leucine, a ketogenic amino acid, also serves to activate the TOR pathway which is one of the most important regulatory pathways governing CO_2_ tolerance (12). Fourth, acetyl-CoA equivalents must also be transported from the mitochondria to other cellular compartments because acetyl-CoA cannot directly cross organelle membranes. We found that the mitochondrial citrate transporter Ymh2 is critical for CO_2_ tolerance likely because it exports citrate from the mitochondrial after which it is converted to acetyl-CoA by ATP-citrate lyase, *ACL1*. The severe infectivity defects of *yhm2*Δ and *acl1*Δ (44) mutant supports the assertion that these processes are also important during infection.

In addition to providing an internally consistent model that explains our genetic and metabolic data, this scheme provides a possible explanation for why some regulatory pathways may be maladaptive during CO_2_ stress. For example, the cAMP-PKA pathway is a well-established mediator of capsule formation (19, 27, 52). The deletion mutant of *PKA1* is essentially devoid of capsule either on the cell surface or secreted into the media, suggesting that activation of the cAMP-PKA pathway is critical to initiation of capsule formation. As previously reported and re-demonstrated in this work, the *pka1*Δ mutant has increased tolerance (12). In this work, we found that deletion of *PKR1*, a negative regulator of the cAMP-PKA pathway, is significantly less fit in CO_2_ (12). The *pkr1*Δ mutant makes a much larger capsule than WT (19). We propose that loss of regulatory pathways such as the cAMP-PKA that function to initiate capsule formation or function early in the pathway, improves fitness in CO_2_ because glucose is not shunted toward capsule production and is available for cellular metabolism. However, the loss of other genes within the capsule biosynthesis process may have either a positive or negative affect on CO_2_ fitness depending on whether they modulate glucose distribution. In this way, capsule-related genes would be present in the GO term sets for those that both increase or decrease CO_2_ fitness (Fig. 1). Further supporting this idea, loss of Uxs1, an enzyme that functions early in the capsule biosynthesis process, leads to increased fitness in CO_2_ (Fig. 4C).

An initially surprising observation was that genes involved in ROS homeostasis were enriched in the set of mutants with reduced CO_2_ fitness. Our subsequent experiments indicate that mitochondrial respiration is increased at host levels of CO_2_. Although there is no increase in the amount of ROS in CO_2_ tolerant H99 cells, our data indicate that this is because multiple ROS compensatory mechanisms maintain ROS homeostasis under these conditions. Loss of glutathione or enzymes in the glutathione biosynthetic pathway leads to both increased ROS and decreased CO_2_ fitness (47). Based on these data, it appears that a key determinant of *C. neoformans* CO_2_ tolerance is the ability of the strain to buffer ROS generated by the increase in mitochondrial activity triggered by host levels of CO_2_. Future studies will be directed to characterizing this capacity in clinical and environmental strains with differences in CO_2_ fitness.

Lastly, we found that the TOR-Ypk1 pathway also contributes to ROS homeostasis both generally and in response to CO_2_ stress. Although this TOR-Ypk1 function is conserved in *S. cerevisiae* (49), it has not been previously demonstrated in *C. neoformans*. In *S. cerevisiae*, the ROS buffering functions of the Tor-Ypk1 pathway are mediated in part by the positive regulation of sphingolipid synthesis (49). Consistent with that conserved function, synthesis of the sphingolipid glucosylceramide is important for both buffering ROS specifically under CO_2_ stress and for fitness at host concentrations of CO_2_. Taken together with previous studies, these data provide a detailed mechanism for how the Tor-Ypk1 pathway governs CO_2_ tolerance through its regulation of lipid homeostasis. As we have previously showed (12), TOR-Ypk1 regulates the remodeling of phospholipid asymmetry by increasing outer membrane phosphatidylserine, a response that likely affects membrane fluidity during CO_2_ stress. Although it is likely that the role of TOR-Ypk1 in sphingolipid biosynthesis mediates CO_2_ tolerance through multiple mechanisms, we show that it is due, at least in part, to the important for buffering mitochondrial ROS induced by host concentrations of CO_2_.

In summary, this work not only provides a near genome-wide profile of the genes required for *C. neoformans* CO_2_ tolerance, but also demonstrates that carbon metabolism, membrane homeostasis and ROS resistance play interconnected roles in establishing that tolerance. More generally, our work demonstrates the extensive remodeling of the cellular physiology that an environmental human pathogen must undergo to adapt to but one aspect of the host environment. The stringency of this process may be one part of the explanation for why a minority of *C. neoformans* strains are able to establish successful infections in mammals.

## Materials and Methods

### Strains and growth conditions

Yeast extract-peptone-2% dextrose (YPD) was prepared according to standard recipes (52). RPMI 1640 without glutamine or sodium bicarbonate was buffered with 165 mM MOPS and pH adjusted to 7. Strains used in this work are described in Table S3. The *C. neoformans* deletion library (2015, 2016, 2020 Madhani plates) was acquired from the Fungal Genetic Stock Center (FGSC). Strains were stored at −80°C in 20% glycerol. Frozen stocks were recovered on solid YPD medium at 30°C for 2 days. To prepare for assays, 3 mL YPD liquid was inoculated per strain and grown overnight, 30°C, shaking at 200 rpm.

### Strain construction

For generation of an independent *yhm2*Δ strain (Table S3), CRISPR/Cas9 short-arm homology (SAH) and transient CRISPR-Cas9 coupled with electroporation (TRACE) methods were used as published (53, 54). CRISPR components were PCR purified according to manufacturer instructions (QIAquick PCR Purification Kit; cat no. 28104) and transformed into H99 via electroporation using the “Pic” setting on a Bio-Rad Micropulser. For gene deletion constructs, *YHM2* SAH Repair 5’ and 3’ oligos designed with a 50 bp sequence matching the flanks of the *YHM2* coding sequence and a 20 bp sequence matching the flanks of the hygromycin (HygB) resistance marker cassette were used to amplify a repair construct, which replaced the entire open reading frame for the target gene with the *HYGB* cassette. Gene deletion constructs were transformed along with Cas9 DNA and two gene-specific sgRNAs into H99. Transformants were selected on YPD medium with 400 µg/mL hygromycin B and knockouts were PCR verified at the 5’ flank with LCR031 and gene specific “5’ KO confirmation primer”, at the 3’ flank with LCR323 and gene specific “3’ KO confirmation primer” and for lack of the native locus with gene specific “orf confirmation” primer pairs (Table S4).

### Flow cytometry-based competitive growth assay

The Madhani mutant collection was screened using a competitive growth assay in which each mutant was compared to the H99 reference strain following previously published procedures (12). Briefly, 96-well plates were prepared with 200 µL YPD per well. Wells were inoculated from growth on solid YPD plates with individual strains to be tested and the reference strain H99-mNeonGreen and grown statically overnight at 30°C. Dilutions were performed to standardize input of unlabeled strains and H99-mNeonGreen in a 1:1 ratio, at 2×10^4^ cells/mL final concentration in RPMI 1640 + 165 mM MOPS, pH 7. Co-cultures were grown at 37°C for 24 hours in ambient air or 5% CO_2_ before analyzing on an Attune NxT Flow Cytometer with CytKick autosampler and Attune Cytometric software. mNeonGreen positive and negative populations were identified by histogram plot with 10,000 single cells counted per sample. The percent of mNeonGreen negative cells in CO_2_ conditions was divided by the percent of mNeonGreen negative cells in ambient conditions to determine the competitive fitness in CO_2_ relative to ambient conditions. Statistical significance was determined by CHI square test in Microsoft Excel. As indicated, additions to the medium were at the following final concentrations; cysteine (200 µM), glutathione (1 mM) and compared mutant cells with supplementation to those without supplementation at 37°C 5% CO_2_.

### Solid agar growth assays

Cells from overnight cultures of indicated strains were washed twice with PBS prior to quantifying OD_600_. Strains were diluted to an OD_600_ :1, followed by ten-fold serial dilutions. 3 µL from each dilution was spotted on agar plates and grown inverted at 37°C in ambient air or at 10% CO_2_ on YPD plates or YPD buffered with HEPES at pH 9. Images were captured at 48hr for growth on standard YPD plates or 144hr for growth on YPD plates at acidic pH.

### Aureobasidin A susceptibility assay

Cells from overnight cultures were washed twice with PBS prior to quantifying concentration on a Countess II FL automated cell counter. Cells were diluted to 1×10^7^ cells/mL to plate on RPMI 1640 + 165mM MOPS pH7 agar plates. A sterile q-tip was saturated in the cell solution before streaking an entire plate to generate a lawn. When plates had dried, a sterile filter paper dot was placed in the center of the plate. Aureobasidin A was added in a 20 µL volume at a final concentration of 100 µg/disk. Plates were incubated at 37°C in ambient air or 5% CO_2_ for 48hr before images were acquired.

### Metabolomic analysis

A 30 mL culture of YPD was inoculated with H99 and grown overnight at 30°C in a 220-rpm shaking incubator. Simultaneously, two groups of 4 flasks containing 50 mL of RPMI media buffered to pH 7.0 with 165 mM MOPS were pre-equilibrated by shaking at 220 rpm in a 37°C incubator with either ambient air or 5% CO_2_ overnight. The following day, late log-phase cultures were back diluted to an OD of 0.5 and allowed to double at 37°C in a 220-rpm shaking incubator. Pre-equilibrated flasks were inoculated to an initial OD of 0.1 from the log phase culture and allowed to grow for 24hr. 100 ODs were collected from each flask, cell pellet rinsed with ice-cold UltraPureTM Distilled Water (Invitrogen), flash frozen in an ethanol dry-ice bath, and stored at −80°C. For extraction, cell pellets were subjected to three rounds of homogenization with glass beads for one minute. Homogenates were resuspended in 2 mL of methanol and transferred to 8 mL glass vials containing 4 mL of cold chloroform (LiChrosolv, Sigma). Samples were then vortexed for 1 min and 2 mL of water (Omnisolv, Sigma) was added. Samples were vortexed for 1 min and then subjected to 10 minutes of centrifugation at 3000 g for phase separation. The aqueous phase was collected (∼3.8 mL) and transferred to a new glass vial. Samples were then frozen at −80°C and submitted to the Harvard Center for Mass Spectrometry (HCMS) where they were dried under nitrogen flow and resuspended in 100 µL of 30% acetonitrile in water.

Untargeted metabolomics was performed using HILIC-UHPLC-MS/MS. A 5 µl injection was separated on a HILICON iHILIC-P Classic column (150 x 2.1mm, 5µm) at 40°C with a ThermoFisher IDX system with HESI source in positive and negative polarity (resolution 120,000; AGC target 1×105; m/z 65-1000). The 45 min LC gradient (0.15 mL/min after 0.5 min. ramp from 0.05 mL/min) used mobile phase A (20 mM ammonium carbonate, 0.1% ammonium hydroxide in water) and B (97% acetonitrile in water): 93% B (0-0.5 min), linear to 40% B (19 min), to 0% B (28-33 min), return to 93% B (36-45 min). Eluent was diverted to waste at 0-0.5 min and 32-46 min. A pooled QC sample underwent AcquireX deepscan (5 levels) in both polarities for MS/MS.

Data were processed in Compound Discoverer 3.2 with the following workflow: (1) MS1 peak detection and integration (threshold-based); (2) retention time alignment; (3) gap-filling by re-integration of low-intensity or noise regions; (4) adduct grouping into single features (neutral monoisotopic mass reported); (5) background subtraction using blanks; (6) median-centered normalization; (7) formula prediction from accurate mass, isotopic pattern, and MS2 (if available); (8) identification via mzVault (local, RT-inclusive) then mzCloud (online MS2); and (9) manual curation of IDs and integrations. One ambient sample exhibited anomalous total intensity and was excluded from differential analysis; normalized areas without this anomalous sample are reported. Peak areas were median-centered and log-transformed prior to statistical analysis, where a Welch corrected two-tailed t-test was performed to compare Ambient (n=3) vs. Elevated (n=4) conditions. Features with log2 (fold-change) >1 and p < 0.05 were considered significantly regulated.

### Mitochondrial membrane potential and reactive oxygen species assays

Cells from overnight cultures of indicated strains were washed twice with PBS prior to quantifying concentration on a Countess II FL automated cell counter. Cells were diluted to 7.5×10^5^ cells/mL in RPMI 1640 + 165mM MOPS pH 7 and aliquoted in 3 ml volumes in 6-well plates, with 3 wells per replicate. Plates were incubated at 37°C in ambient air or 5% CO_2_ for 24 hr. At 24 hr, wells were combined, concentration was quantified by cell counter and 1 × 10^6^ cells were aliquoted into a 24-well plate per staining condition. Cells were resuspended in 1 ml RPMI containing 2.5 µM JC-1 (catalog no. M34152; Thermo Fisher) or 16 µM DCFDA (cagalog no. D6883; Sigma) and incubated at 37°C in ambient air or 5% CO_2_ for 1 hr. Cells were washed once with PBS, then resuspended in 500 µl PBS and analyzed on an Attune NxT Flow Cytometer with Attune Cytometric software. JC-1 staining was analyzed by dividing the mean fluorescence intensity (MFI) resulting from excitation with the blue (488nm) laser in the BL2 emission filter (574/26) by the MFI in the BL1 emission filter (530/30), i.e. red/green ratio of single yeast cells. DCFDA staining was analyzed by measuring the MFI of single yeast cells in the BL1 emission filter (530/30) following excitation with the blue (488nm) laser.

### Quantification of NAD/NADH and NADP+/NADPH ratios

Cells from overnight cultures of H99 were washed twice with PBS prior to quantifying concentration on a Countess II FL automated cell counter. Cells were diluted to 7.5×10^5^ cells/mL in RPMI 1640 + 165 mM MOPS pH 7 and aliquoted in 3ml volumes in 6-well plates, with 3 wells per replicate, performed in biological triplicate. Plates were incubated at 37°C in ambient air or 5% CO_2_ for 24 hours. At 24hr, replicate wells were combined, concentration was quantified by automated cell counter and two tubes of 1 × 10^7^ cells were pelleted per replicate/condition to allow simultaneous analysis of the oxidized and reduced form from each replicate. Cells were frozen at −80°C and lyophilized overnight. Manufacturer kit instructions were followed for NAD+/NADH (BioAssay E2ND-100) or NADP+/NADPH (BioAssay Systems E2NP-100) quantification.

### Mouse model of cryptococcosis

This study was performed according to the guidelines of NIH and the University of Georgia Institutional Animal Care and Use Committee (IACUC). The animal models and procedures used have been approved by the IACUC (AUP protocol number: A2023 03-033-A3). CD1 female mice (022CD1) with a body weight of 18-25 grams were purchased from Charles River. The mice were housed with a max of 5 per cage with a 7am-7pm 12hr dark/12hr light cycle. They were fed LabDiet 5053 irradiated Picolab Rodent Diet 20. The water was provided by the automated watering system connected to the rack, which has several filtration systems. The ambient temperature was between 20-24°C with 30-70% humidity. For intravenous inoculation (IV), wild-type and mutant cells were mixed in a 1:1 ratio prior to inoculation with 1 × 10^5^ colony forming units (CFUs)/animal in 100 µl volume. Organs were harvested at day 4 and homogenized in 2 mL of cold sterile PBS using an IKA-T18 homogenizer. Homogenized organs were serially diluted, plated onto YNB and YNB with 400 µg/mL hygromycin and incubated at 30°C for 2 days before counting CFUs. Neither the *acl1*Δ nor *yhm2*Δ strains were detected in the brain. For intranasal inoculation (IN), wild-type and mutant cells were mixed in a 1:1 ratio prior to inoculation with 5 × 10^4^ CFU/animal in a 50 µl volume. Fungal burden quantification was performed as above.

### Western blotting

Overnight cultures of H99 were rinsed in water and resuspended to spot 5 × 10^7^ cells in 20 µl in biological triplicate on an RPMI 1640 + 165 mM MOPS pH7 agar plate. Replicate plates were incubated at 37°C overnight. Prior to CO_2_ exposure, a 0-hour time point was collected, then one plate was shifted to 37°C + 5% CO_2_, while one remained at 37°C ambient CO_2_ for 2 hours. At 2 hr, all replicates were swabbed from the plates into individual tubes containing 500 µl 0.2N NaOH. Suspensions were incubated at room temperature for five minutes. Cells were pelleted and resuspended in 20 µl reducing sample buffer. Samples were boiled for 5 minutes, then pelleted at 5,000rpm for 3 min before removing supernatant to fresh tubes. 10 µl from each replicate was loaded into duplicate 10% SDS-PAGE gels and run at 80V. Samples were transferred to a nitrocellulose membrane for 1 hour at 100V. Membranes were blocked with 5% bovine serum albumin (BSA) in tris-buffered saline with 0.1% tween 20 (TBST) for 1 hr at RT, then incubated with 1:10,000 rabbit anti-Hog1 (generous gift from Yong-Sun Bahn) or rabbit anti-phospho-p38 MAPK (catalog no. 4092S; Cell Signaling Technology) @ 1:10,000 in 5% BSA/TBST overnight at 4°C. Membranes were washed 3 times for 5 min with TBST, then incubated for 1 hr at RT with 1:10,000 goat anti-rabbit horseradish peroxidase (HRP) (catalog no. STAR208P; Bio-Rad) in 5% BSA/TBST. Membranes were washed 3 times for 5 min with TBST, then developed with chemiluminescent substrate (catalog no. 1705060; BioRad) and imaged on a myECL imager (ThermoFisher).

### Statistical analyses

Data were graphed and analyzed in GraphPad Prism 9 software or Microsoft Excel. Statistical analysis and graph descriptions, including P values are provided in figure legends.

## Acknowledgements

We acknowledge the support of grant R01AI147541 from the National Institute of Allergy and Infectious Disease (DJK and XL) and the University of Georgia Gene E. Michaels Fund (XL). We acknowledge the Madhani Lab (UCSF) and the National Institutes of Health funding (R01AI100272) for the creation of the *Cryptococcus neoformans* deletion mutant collection resource used in this work. We thank Andy Alspaugh (Duke) for providing *RIM101* mutant strains. We thank Yun-Sun Bahn (Yonsei University) for providing samples of anti-*Cryptococcus neoformans* Hog1.

